# FALCONET: an R package to accelerate automatic visualisation of genome scale metabolic models

**DOI:** 10.1101/662056

**Authors:** Hongzhong Lu, Zhengming Zhu, Eduard J Kerkhoven, Jens Nielsen

## Abstract

**Summary:** FALCONET (FAst visuaLisation of COmputational NETworks) enables the automatic formation and visualisation of metabolic maps from genome-scale models with R and CellDesigner, readily facilitating the visualisation of multi-layers omics datasets in the context of metabolic networks.

**Motivation:** Until now, numerous GEMs have been reconstructed and used as scaffolds to conduct integrative omics analysis and *in silico* strain design. Due to the large network size of GEMs, it is challenging to produce and visualize these networks as metabolic maps for further in-depth analyses.

**Results:** Here, we presented the R package - FALCONET, which facilitates drawing and visualizing metabolic maps in an automatic manner. This package will benefit the research community by allowing a wider use of GEMs in systems biology.

**Availability and implementation:** FALCONET is available on https://github.com/SysBioChalmers/FALCONET and released under the MIT License.

**Supplementary information:** Supplementary data are available online.

## 1 Introduction

In the data-driven era, genome scale metabolic models (GEMs) have become useful tools for the integrative analysis of large omics datasets. High-quality visual maps of these metabolic networks can greatly aid researchers to appreciate and interpret the complex interactions among reactions, genes and proteins; e.g. directly visualizing changes in metabolic activities. Moreover, high-quality metabolic maps are beneficial in metabolic engineering, to visualize rational design strategies of cell factories by identifying *in silico* gene targets related to high yield of bioproducts. Currently, there are various software solutions supporting visualisation of metabolic models, like Omics (Droste, et al., 2011), CellDesigner (Funahashi, et al., 2008) and Escher (King, et al., 2015). However, these tools cannot automatically generate metabolic maps, and drawing a high-quality metabolic map for a non-model organism can take several months. Consequentially, there is a strong demand to be able to produce metabolic maps using a flexible and automatic approach. Due to the complex connections in metabolic networks, layout adjustment is another major challenge during the automatic formation of metabolic maps. In addition, the metabolic maps produced using the automatic methods should be able to be easily used in different platforms after a few format curations. While some tools, like Metdraw (Jensen and Papin, 2014), can produce metabolic maps in batch, the export format of these tools cannot be reused for the above popular visualisation tools. It is therefore imperative that a novel tool is required to overcome all these bottlenecks.

Here we present FALCONET – an R package to accelerate the formation and visualisation of GEMs on a large scale. It not only enables the automatic visualisation of GEMs in html format using R, it can furthermore produce metabolic maps of high quality in batch by leveraging CellDe-signer (Funahashi, et al., 2008). Moreover, FALCONET supports direct visualisation of multi-layers omics through the reconstructed map. Maps generated in xml format can readily be used in other platforms through some essential format curation.

## 2 Implementation details

### 2.1 Choosing interesting reactions sets

As the input for FALCONET, the output of a metabolic model with com-munity-standard reaction and metabolite annotations from COBRA (Schellenberger, et al., 2011) and RAVEN (Agren, et al., 2013) in Excel or CSV format should be provided. In FALCONET, reactions can alternatively be filtered based on subsystems (Fig.1A). Meanwhile, functionality is provided to specifically select those reactions that are connected to specific genes and/or metabolites.

**Fig 1.**
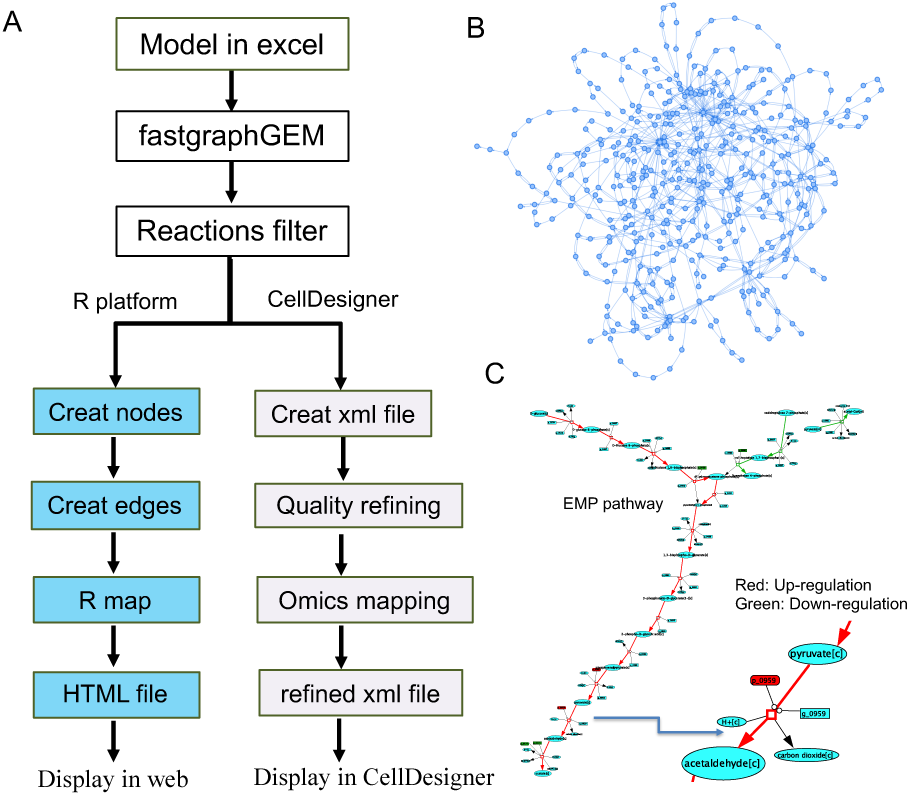
FALCONET makes it possible to produe metabolic maps in batch with aid of CellDesigner. (A) Main procedueres to prepare the maps in FALCONET. (B) Map used in the R platform for a medium size of yeast metabolic model, which contains 270 reactions. (C) An example of displaying multi-layer omics data on the metabolic map in xml format. The visualisation is realized by CellDesigner.

### 2.2 Draft maps

Firstly, the FALCONET package could produce xml files used for CellDesigner. In xml file, the coordinates of the metabolites, genes and proteins within the map were defined automatically. Subsequently, the connections among the metabolites determined by reactions were added in the map. Next, a general gene (protein) symbol is connected with each reaction. Sometimes a metabolic model contains different compartments, thus the related transport reactions in the model were found and added automatically to connect the metabolites from different compartments in the same subsystem. For the map displayed in R platform (Fig.1B), only the metabolites and reaction ID were shown. For each reaction, the reactant and product are connected through the reaction ID. More visualisation example using R platform are available as Supplementary file.

### 2.3 Refining maps for omics visualisation

The draft maps in xml format need to be imported into CellDesigner for automatic layout adjustment. Next, the map quality can be improved through adjusting the size and color of nodes defined in the map generated by FALCONET. The refined maps can be used to readily visualize proteomics, transcriptomics and fluxomic. Fluxomic can be directly mapped based on reaction ID, while the gene-protein-reaction relations (GPR) can be used to map proteomics and transcriptomics data. If a reaction contains several isoenzymes, the mean values of gene expression is used to reflect the changes. By contrast, if a reaction is catalyzed by a complex, the minimum expression value of all subunits is used.

### 2.4 Export

FALCONET provides two options for visualisation of metabolic maps (Supplementary file). For direct visualisation used in the R platform, it can export the output as an html file. Secondly it can produce a refined xml files mainly to be used for CellDesigner. Existing tools, like Escher-Converter (https://github.com/draeger-lab/EscherConverter), can trans-form the xml file into an SVG file or JSON file format to facilitate reuse of the maps with other widely-used tools, such as Omics and Escher.

## 3 Usage scenario

As a typical example of using FALCONET, the subsystem map for purine metabolism of *Saccharomyces cerevisiae* (https://github.com/SysBioChalmers/yeast-GEM) was drawn (Supplemen-tary file) rapidly in comparison to fully manual efforts. Additional usage scenarios are displayed in Supplementary file.

As an example of visualizing multi-layers omics data with FALCONET, physiological and omics data from *S. cerevisiae* chemostat cultivations under 30 °C and 38°C were used. Flux distributions were obtained from yeast GEM using pFBA (Lewis, et al., 2010). Shown in in Fig.1C is the EMP pathway for *S. cerevisiae* as generated automatically with FALCONET, including automatic layout adjustment with CellDesigner. Here only the reactions carrying fluxes were shown. Using the GPR relations, the transcriptomics and the proteomics data were mapped onto the corresponding gene and protein identifiers for each reaction. The increased expression of genes or proteins for the related reactions were marked as red. The flux values with significant up and down-regulated tendencies were visualized using red and green color, respectively. The functions contained in FALCONET therewith make it possible for users to extract import clues for metabolic regulation under specific genotypes or external environments.

