## Supplementary file for "FALCONET: an R package to accelerate automatic visualisation of genome scale metabolic models"

### Usage examples of FALCONET

#### 1 More detailed usage cases of FALCONET in CellDesigner platform

##### 1.1 Example metabolic map for subsystem

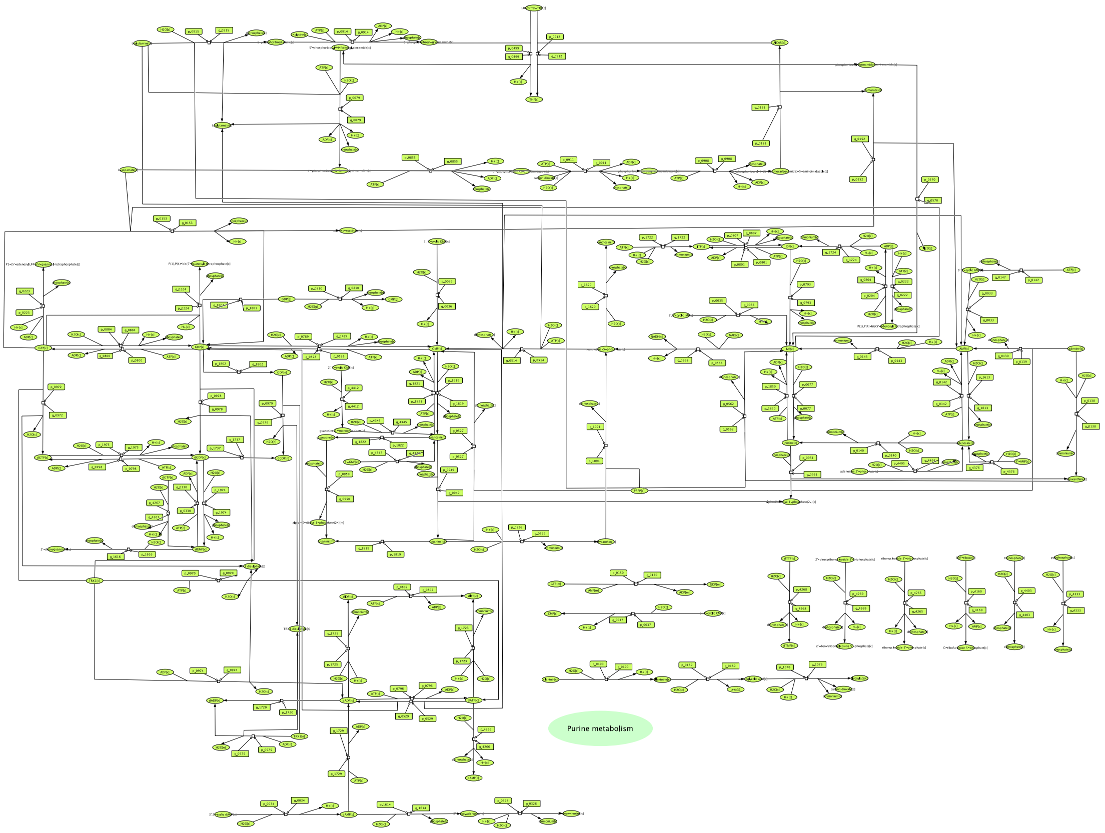


Figure S1 Map for 'purine metabolism' from yeast GEM (version 8.3) drawn by FALCONET and CellDesigner 4.4. After the automatic layout adjustment, some manual correction was done to refine the map quality.

##### 1.2 Find the reactions connected with specific metabolites

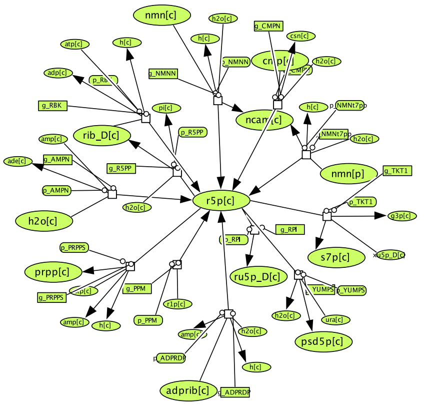


Figure S2 All reactions connected with r5p from *E. coli* iML1515. With FALCONET, all the reactions connected with specific metabolites could be chose to produce the xml file, which can help understand the systematic roles of the target metabolite in the network.

##### 1.3 Find the reactions connected with specific genes

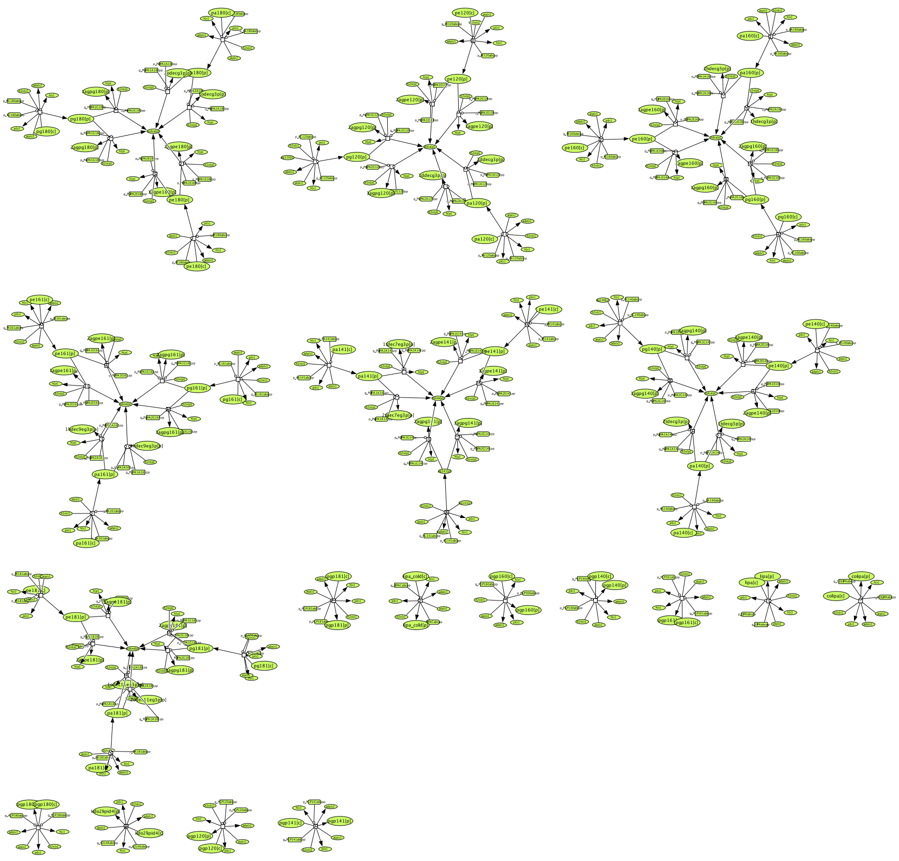

Figure S3 All reactions connected with genes - ‘b0914 or b3821’ from *E. coli* iML1515. With FALCONET, all the reactions connected with specific genes could be chose to produce the xml file, which can help understand the systematic roles of the target genes in the network.

##### 1.4 Choose the reactions carrying flux from specific subsystems

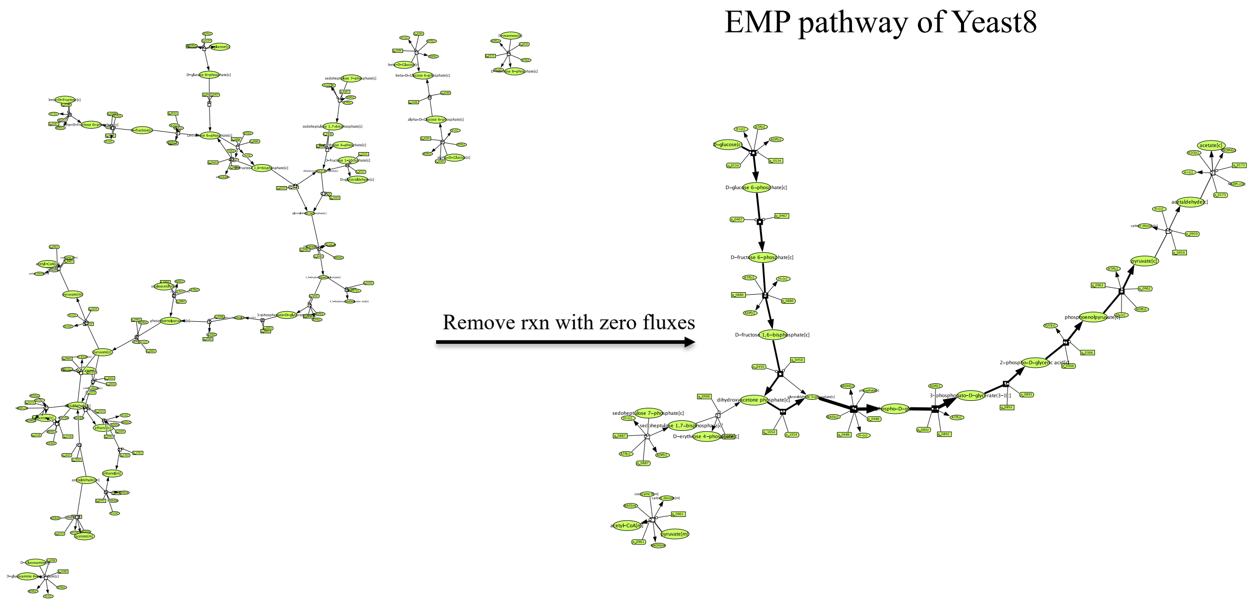


Figure S4 FALCONET could produce the maps only for those reactions which carry fluxes. As the original map can be very complex due to the large size, the refined map only contained reactions carrying fluxes could contribute to an easy visualization.

#### 1.6 Parameters setting in the CellDesigner for the automatic layout adjustment

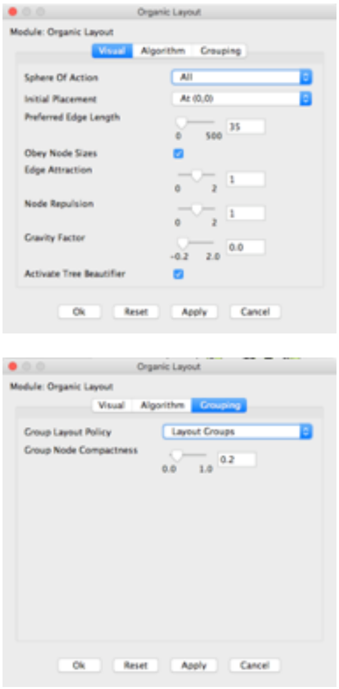


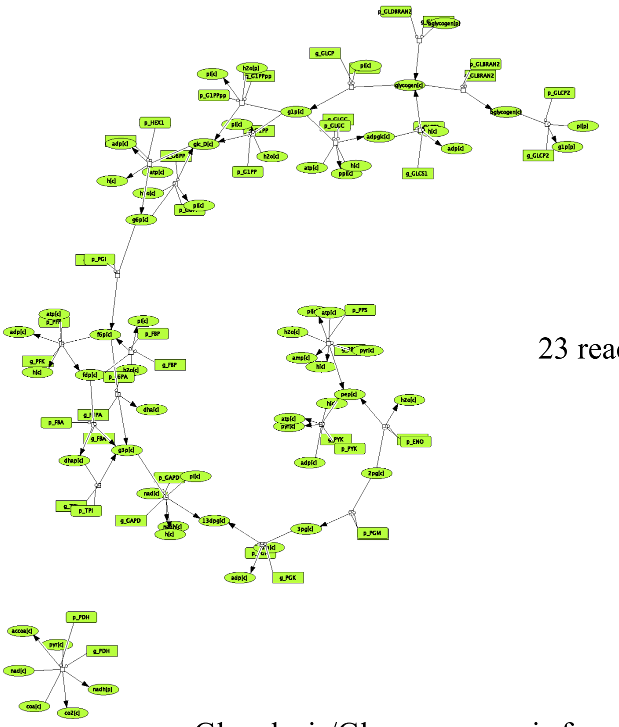

23 reactions

for subsystem with <100 rxns

Glycolysis/Gluconeogenesis from iML1515

Figure S5 Parameters setting for the organic layout adjustment in CellDesigner 4.4. Using the tuned parameters provided by CellDesigner 4.4, a map of high quality with better layout could be obtained.

#### 2 More detailed usage cases of FALCONET in visualization of maps in R platform

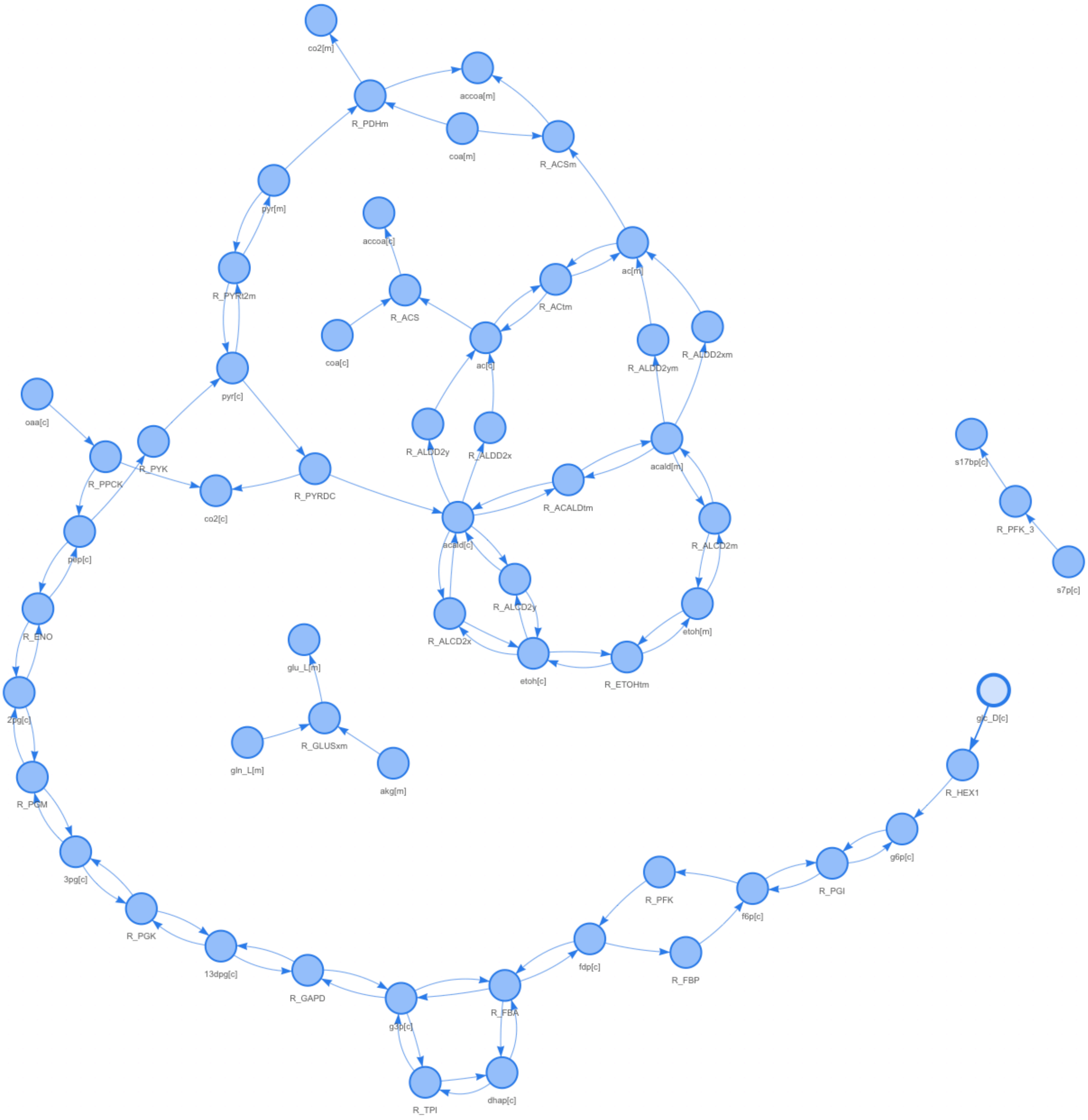


Figure S6 Visualization of metabolic maps based on R platform. The FALCONET could produce the map with html file as an output which can be visualized using the web. An easy zoom-in and zoom-out of the map in html format can be realized for an easy visualization.

#### 3 Export of FALCONET

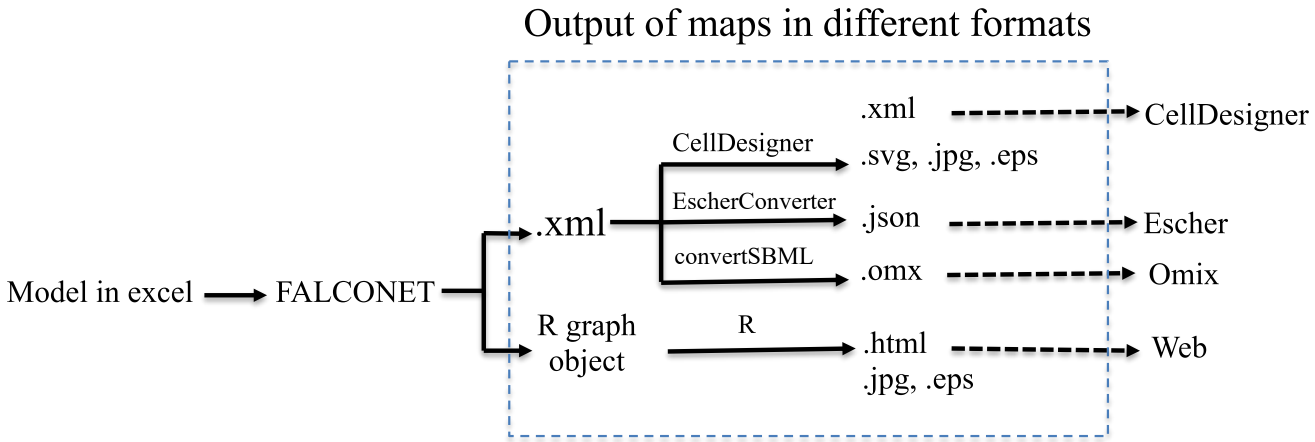


Figure S7 The maps in xml format could be used in Escher and Omics platform after a few curation in the format.
